# Ankle muscles drive mediolateral center of pressure control to ensure stable steady state gait

**DOI:** 10.1101/2021.03.31.437904

**Authors:** A.M. van Leeuwen, J.H. van Dieën, A. Daffertshofer, S.M. Bruijn

## Abstract

During steady-state walking, mediolateral gait stability can be maintained by controlling the center of pressure (CoP). The CoP modulates the moment of the ground reaction force, which brakes and reverses movement of the center of mass (CoM) towards the lateral border of the base of support. In addition to foot placement, ankle moments serve to control the CoP. We hypothesized that, during steady-state walking, single stance ankle moments establish a CoP shift to correct for errors in foot placement. We expected ankle muscle activity to be associated with this complementary CoP shift. During treadmill walking, full-body kinematics, ground reaction forces and electromyography were recorded in thirty healthy participants. We found a negative relationship between preceding foot placement error and CoP displacement during single stance; steps that were too medial were compensated for by a lateral CoP shift and vice versa, steps that were too lateral were compensated for by a medial CoP shift. Peroneus longus, soleus and tibialis anterior activity correlated with these CoP shifts. As such, we identified an (active) ankle strategy during steady-state walking. As expected, absolute explained CoP variance by foot placement error decreased when walking with shoes constraining ankle moments. Yet, contrary to our expectations that ankle moment control would compensate for constrained foot placement, the absolute explained CoP variance by foot placement error did not increase when foot placement was constrained. We argue that this lack of compensation reflects the interdependent nature of ankle moment and foot placement control. We suggest that single stance ankle moments do not only compensate for preceding foot placement errors, but also assist control of the subsequent foot placement. Foot placement and ankle moment control are ‘caught’ in a circular relationship, in which constraints imposed on one will also influence the other.

## Introduction

Muscle activity is required to coordinate the base of support (BoS) with respect to the center of mass (CoM) in humans in response to perturbations and during steady-state walking ^1,2^. Proper coordination between the BoS and CoM ensures gait stability ^3-5^. The most dominant strategy to achieve this appears to be the control of foot placement ^1,4^. Accurate foot placement positions the center of pressure (CoP) such that the ground reaction force generates an adequate moment to accelerate the CoM away from the lateral border of the BoS ^4,5^, through which falls can be prevented.

Foot placement in accordance with stability constraints may not always be possible due to environmental or individual constraints. A missing tile in the pavement may force one to step elsewhere. Motor (and/ or sensory) noise and insufficient force generating capacity may also prevent stepping to a proper location. Stroke patients with a high fall risk show poor coordination between the CoM and foot placement ^6^. In a previous study, we “impaired” coordination of foot placement relative to the CoM by instructing healthy subjects to place their feet on projections on the treadmill at a fixed step width. This foot placement constraint turned out effective in preventing participants to vary their foot placement according to variations in their CoM state. Relying less on foot placement control proved possible for healthy subjects, although it may demand a compensatory strategy to maintain stability.

A potential compensatory strategy is the so-called mediolateral ankle strategy ^7,8^. It aims to shift the CoP relative to the CoM during (single) stance. As such, the coordination between the BoS and the CoM can be improved before the next foot placement. An interaction between foot placement and execution of the ankle strategy has been found after perturbations ^7,9,10^ and has also been demonstrated during (unconstrained) steady-state gait ^11^. For instance, walking with constrained ankle moments led to a compensatory wider step width ^12^. This need for compensation when ankle moments are limited suggests that during unconstrained steady-state gait, foot placement control is complemented by ankle moment control. A serial execution of the ankle and foot placement strategies allows for an early response to a perturbation by ankle moments, followed by a larger response (in terms of CoP displacement) through foot placement ^7^. Conversely, inaccurate foot placement can be accommodated for by subsequent ankle moment control ^8^.

We employed the foot placement model proposed by Wang and Srinivasan ^13^ to reflect an (active) control strategy for foot placement. In previous work we used the relative explained variance of this model to analyze the degree of foot placement control. Here, we focused on the residual variance of the same model, as an error in foot placement. This error (unexplained variance) can be attributed to mere motor noise or relaxed control.

If detrimental to stability, this error in foot placement may potentially be compensated for by ankle moment control. To identify an (active) mediolateral ankle strategy during steady-state gait, we correlated the error term of the foot placement model with the subsequent CoP shift underneath the stance foot.

We hypothesized that in steady-state walking, the mediolateral CoP displacement during single stance can be predicted by the error between predicted and actual foot placement (H1a). Furthermore, we hypothesized that muscles around the ankle actively control the CoP shift (H1b). To test this, we correlated soleus, tibialis anterior and peroneus longus activity, to mediolateral CoP displacement during single stance.

In seeking additional support for the execution of the mediolateral ankle strategy during steady-state walking, our participants wore a shoe with a narrow ridge underneath the sole (LesSchuh). With this constraint we intended to limit the variation in ankle moments and – by this – to test whether this limitation leads to a decreased absolute explained variance when predicting mediolateral CoP displacement based on the foot placement error (H2).

In addition, we sought to assess the compensatory potential of the ankle strategy. In this context, we hypothesized that diminishing foot placement control leads to an increase of the absolute explained variance by the ankle strategy model (H3a). Furthermore, We expected a similar effect for the relation between ankle muscle activity and CoP shifts, reflecting an increased active contribution of this control strategy (H3b).

Finally, we investigated the role of gait speed on the contribution of ankle moment control to mediolateral gait stability. Tight foot placement control is arguably more important at normal than at slow walking speed ^14^. By the same token, foot placement responses to vestibular perturbations are larger at a higher than at a lower stepping frequency ^15^, while for the execution of the ankle strategy, the opposite seems to be the case ^15^. Perhaps, the time duration of single stance determines the efficacy of the ankle strategy. A longer single stance duration provides more time during which an ankle moment can be applied and allows for a larger center of pressure shift. This suggests a larger ankle strategy contribution during slow as compared to normal steady-state walking. Along these lines, we hypothesized the contribution of ankle moment control to be higher at a slow walking speed, as reflected by a higher absolute explained variance given by the relation between CoP shifts and foot placement error (H4a). In accordance, we expected the absolute explained variance for the relation between ankle muscle activity and CoP shifts to be higher at slow speed as well (H4b).

In summary, this study focuses on whether inaccurate foot placement is compensated for by (active) ankle moment control. To this end, we investigated correlations between foot placement errors, center of pressure shifts and preceding ankle muscle activity. Furthermore, we assessed the effects of ankle moment and foot placement constraints on these relationships. Finally, we tested whether walking speed determines the contribution of (active) ankle moment control during steady-state walking.

By identifying active ankle moment control during steady-state walking we contribute to the understanding of neural control of mediolateral gait stability. Understanding which control strategies contribute (actively) to a stable gait pattern can help to understand and/or intervene upon decreased gait stability in elderly, pathological or prosthetic gait and may contribute to the control of walking exoskeletons.

This study’s preregistered hypotheses, protocol and sampling plan can be found on OSF: https://osf.io/74pn5. The data can be found at: https://doi.org/10.5281/zenodo.4229851^16^. In addition, the code for the analysis of this study can be found at: https://surfdrive.surf.nl/files/index.php/s/d1uV32TEPrirR23 (and will be published on Zenodo when accepted for publication).

## Methods

Most of the methods have been described before in van Leeuwen, et al. ^12^, overlapping the current methods section.

### Participants

As previously described ^12^, only participants capable of walking for a longer duration (± 60 minutes in total) were included. Participants reporting previous or ongoing sports injuries or other motor impairments, which could affect their gait pattern, were excluded. Participants suffering from balance issues (self-reported) were also excluded. An initial sample of ten participants was included as we used a Bayesian sequential sampling approach. Subsequently, recruitment of participants continued until a threshold of meaningful evidence was reached ^17^. We indicated this threshold by means of a Bayes Factor (BF) ^17^. We set the threshold to a BF10 or BF01 of 10 or 0.1 (indicative of strong relative evidence) for either the null or the alternative hypothesis (based on our main outcome measures). Since not all outcome variables reached the threshold for BF10 or BF01, we continued recruitment until the pre-determined maximum of 30 participants, after accounting for drop-out, was attained.

35 healthy participants completed the experiment according to the instructions. The data of four participants were discarded because of technical malfunctioning of equipment, and data of one participant were excluded in view of pronounced toeing out in his ordinary gait pattern. Ultimately, 30 participants (aged 30 ± 7 yrs, 70 ± 13 kg, 1.73 ± 0.08 m (mean ± sd)) were included in the data analysis.

Prior to participation, participants signed an informed consent. Ethical approval (VCWE-2018-159) had been granted by the ethics review board of the faculty of Behavioural and Movement Sciences at the Vrije Universiteit Amsterdam. The experiments were performed in accordance with relevant guidelines and regulations.

### Procedure

Participants were asked to walk on a treadmill during three conditions at normal (1.25 · √(leg length) m/s) and slow (0.63 · √(leg length) m/s) walking speeds, normalized to leg length ^18^. Stride frequency was controlled by means of a metronome. Participants were asked to time their right heel strikes to the metronome beat. The imposed frequency was set to the average preferred stride frequency as demonstrated during the final 100 steps of a familiarization trial for each speed.

**Table 1.**
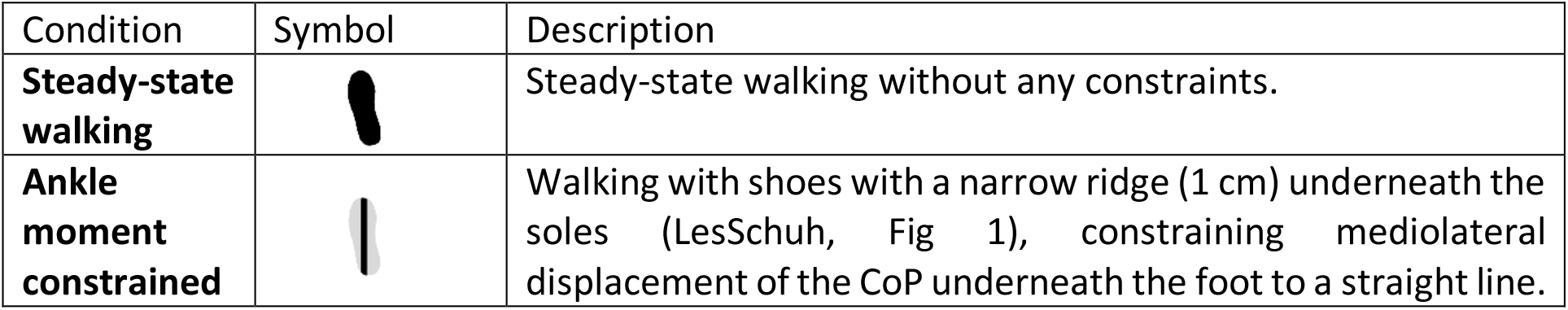
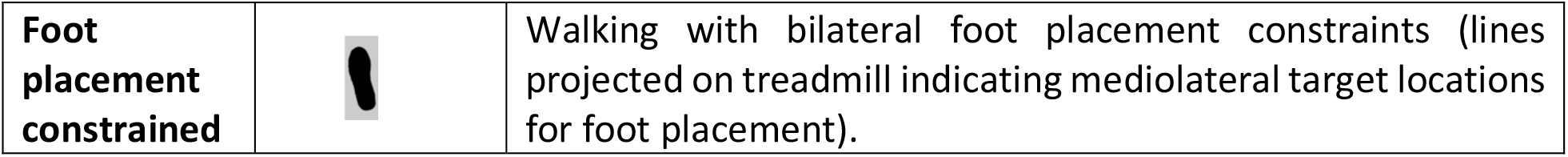
Conditions performed at normal and slow walking speeds. As published in ^12^.

| Condition | Symbol | Description |
| --- | --- | --- |
| **Steady-state walking** |  | Steady-state walking without any constraints. |
| **Ankle moment constrained** |  | Walking with shoes with a narrow ridge (1 cm) underneath the soles (LesSchuh, Fig 1), constraining mediolateral displacement of the CoP underneath the foot to a straight line. |
| **Foot placement constrained** |  | Walking with bilateral foot placement constraints (lines projected on treadmill indicating mediolateral target locations for foot placement). |

The order of the conditions was randomized across participants and speeds. Before commencing the experiment, participants performed a five-minutes (two minutes at normal walking speed, three minutes at slow walking speed) treadmill familiarization trial without any constraints imposed. Additionally, participants practiced walking in LesSchuh on the treadmill prior to the data collection to ensure they were familiar with a constrained ankle moment. To ensure that all trials contained at least 200 consecutive strides, trials at normal walking speed lasted five minutes and trials at slow walking speed lasted ten minutes. Between trials, sufficient breaks were provided to prevent fatigue as verified by subjective report.

### Foot placement constraint

Beams were projected on the treadmill constraining mediolateral foot placement to a fixed step width. Average step width was derived from the final 100 steps of the familiarization trial, based on the CoP recorded by the instrumented treadmill. This average step width was imposed by projecting beams on the treadmill and participants were instructed to place their foot in the middle of the beam. This beam became visible following toe-off, triggered by a force threshold, to prevent modification of the CoM swing phase trajectory by compensatory push-off modulation ^19^.

### Ankle moment constraint

A shoe (LesSchuh) with a narrow (1 cm) ridge attached to the sole, as a limited base of support, was used to avoid a mediolateral shift of the CoP underneath the stance foot. LesSchuh was designed to limit ankle moments. At the same time, it still allows for anteroposterior roll-off and subsequent push-off because the material of the ridge bends with the sole in anterior-posterior direction. Participants were asked to walk on the ridge, without touching the ground with the sides of the shoe’s sole. Participants were also instructed to place their feet in a similar orientation as they would without the constraint, to avoid a “toeing-out strategy” ^20^. Toeing-out could induce a mediolateral shift of the center of pressure after foot placement despite the narrow BoS.

### Data collection

#### Force plate

Participants walked on an instrumented dual-belt treadmill. Ground-reaction forces and moments were obtained from the force plates embedded in the treadmill at 1000 samples/s. From this data the CoP was calculated. The treadmill was calibrated using an instrumented calibration pole ^21^.

#### Kinematics

Full-body kinematics were measured using two Optotrak (Northern Digital Inc, Waterloo Ont, Canada) cameras directed at the center of the treadmill, sampling at a rate of 50 samples/s. Cluster markers, containing 3 single markers, were attached to the feet, shanks, thighs, pelvis, trunk, upper arms and forearms, and anatomical landmarks were digitized using a six-marker probe.

#### Electromyography

Bipolar surface electromyography (EMG) served to measure bilateral muscle activity from 16 muscles expected to contribute to gait stability. For the current study, we focused on m. peroneus longus (PL), the m. soleus (SO), and m. tibialis anterior (TA) muscles, hypothesized to contribute to the execution of a mediolateral ankle strategy (for other muscles recorded, we refer to the preregistered protocol: https://osf.io/74pn5). A 16-channel Porti EMG device (TMSi, Enschede, Netherlands) was used to record the three muscles bilaterally. Data were sampled at a rate of 2000 Hz. Surface EMG electrodes with a diameter of 22 mm were positioned on the skin conform SENIAM guidelines ^22^.

### Data Analysis

For all subjects and trials, we analyzed 200 consecutive strides. These were the final 200 strides of each trial unless data quality (better marker visibility, less noise) urged selection of earlier strides.

### Gait event detection

In the offline analysis for all conditions, gait events (heel strikes & toe-offs) were detected based on the characteristic “butterfly pattern” of the center of pressure as derived from force plate data ^23^. A step was defined as the period between toe-off and heel strike. Mid-swing was defined at 50% of the step.

### Center of mass

For every segment an estimation of the segment’s mass was made based on linear regression including the segment’s circumference and length as predictors and regression coefficients based on gender. The CoM was expressed as a percentage of the longitudinal axis of the segment ^24,25^. The total body CoM was derived from a weighted sum of the body segment’s CoMs. The derivative of the mediolateral position of the CoM (CoM_pos_) was taken to obtain the mediolateral velocity of the CoM (CoM_vel_).

### Center of pressure

Force plate data were aligned with the Optotrak coordinate system to be able to express the CoP in the local coordinate systems of the feet. This local coordinate system (Fig 2) was defined based on the malleoli, tip of the second toe and the calcaneus as bony landmarks. The ankle joint center was defined as the midpoint between the malleoli. A temporary forward axis was defined by the vector from the calcaneus to the toe tip. The cross product between the vector from the calcaneus towards the ankle joint center and the temporary forward axis computed the horizontal (mediolateral) axis. The cross product between the temporary forward axis and the horizontal axis computed the vertical axis. Lastly the forward axis was computed by the cross product of the horizontal and vertical axis. In our analysis we focused on CoP shifts along the local mediolateral axis.

**Fig 1.**
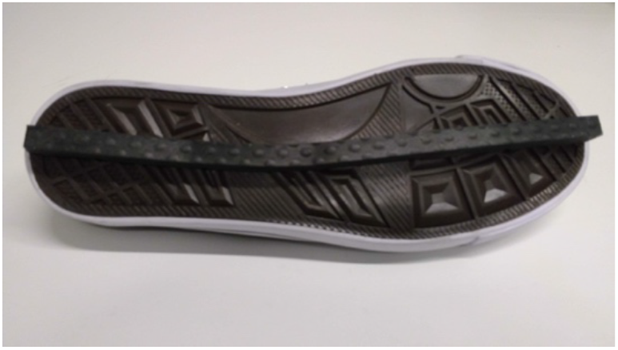
LesSchuh. Shoe with ankle moment constraint (width of the ridge is 1 cm). Figure as published in ^12^.

**Fig 2.**
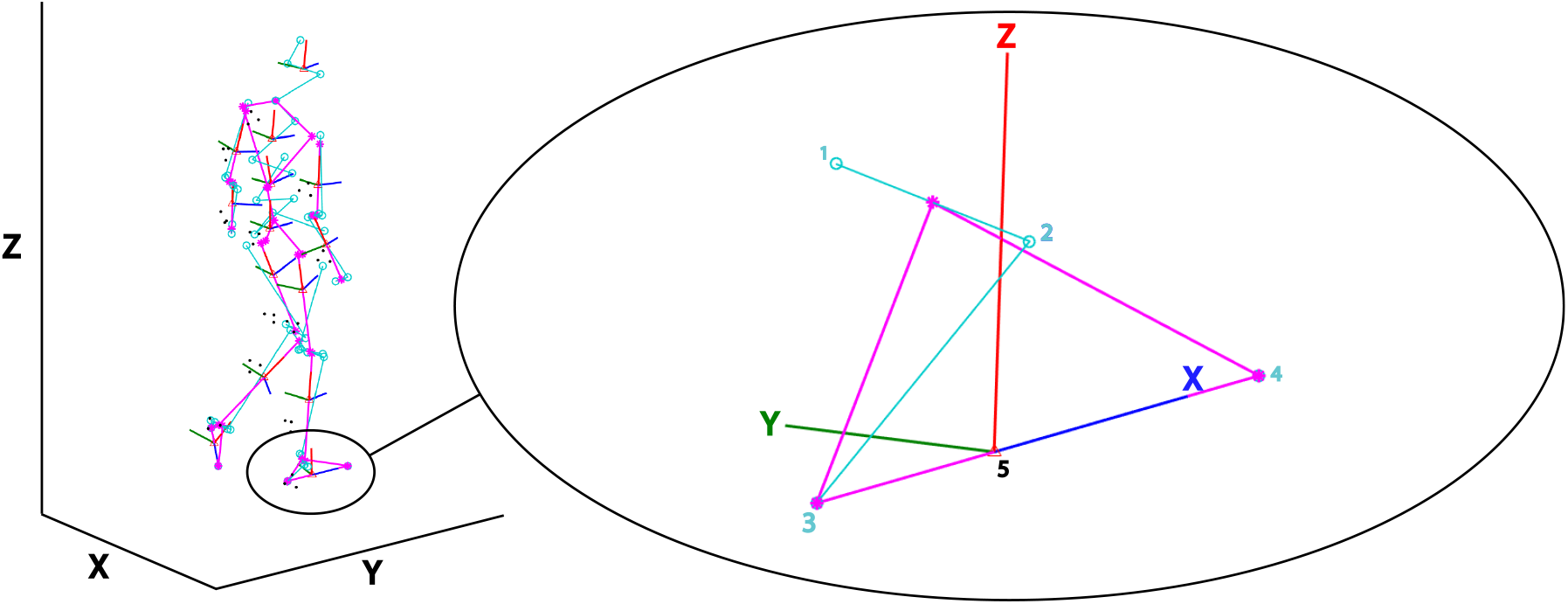
Local foot coordinate system. Based on digitized anatomical landmarks a model of this example participant was constructed. The full model has been depicted on the left. Magnified is the local coordinate system (X,Y,Z) of the right foot with its anatomical landmarks (light blue circles). The local coordinate system was constructed based on the medial (1) and lateral (2) malleoli, the calcaneus (3) and the second toe tip (4). The origin of the constructed coordinate system lies at the estimated foot’s CoM position (5). The red, dark blue and green lines represent respectively the local vertical (Z), forward (X) and mediolateral axes (Y). We defined our mediolateral CoP shift along the anatomical Y-axis in green. Note that in this figure this mediolateral axis points positively to medial, yet in our analysis we flipped the sign to define a lateral shift as positive and a medial shift as negative. This figure was created using Matlab 2021a (https://www.mathworks.com/products/matlab.html) and Adobe Illustrator CC 2018 (https://www.adobe.com/nl/products/illustrator.html).

### EMG processing

EMG data were high-pass filtered at 20 Hz, rectified, and low-pass filtered at 50 Hz following Rankin, et al. ^1^. Every stride was time normalized to 1000 samples.

### Multiple linear regression models

For the foot placement model (Model 1), multiple linear regression analysis was performed with mediolateral foot placement (FP) as the dependent variable and CoM state (position: pos, velocity: vel) variables ([CoM_pos_ CoM_vel_]), as the independent variables. The Matlab code for this has been separately published on Github ^26^. FP was determined at mid-stance, expressed with respect to the position of the contralateral foot and demeaned. Moreover, swing phase CoM_pos_ and CoM_vel_ were expressed with respect to the position of the stance foot and demeaned. The predictors’ timeseries were time normalized to 51 samples for every step (from toe-off to heel strike).

For the ankle strategy model (Model 2), we performed a multiple linear regression analysis with the mediolateral CoP displacement (CoPshift) as the dependent variable and the foot placement error as the independent variable. CoPshift was defined as the demeaned CoP shift in the local coordinate system of the foot (This is a deviation from the preregistered plans. See “Deviations from the preregistered plans”, I). We quantified this CoP shift as the average mediolateral shift during single stance with respect to the initial CoP position at contralateral toe-off ^7^. Error_fp_ denoted the residual between actual foot placement and foot placement predicted by CoM state (model (1)) at respectively mid-swing and heel strike. Foot placement (fp) errors represented steps that were either too lateral (positively signed errors) or too medial (negatively-signed errors). Error_as_ denotes the difference between the actual CoP_shift_ and its prediction.

In the muscle model (Model 3), mediolateral CoP displacement (CoPshift) was the dependent variable and soleus, tibialis anterior and peroneus longus EMG amplitudes [EMG_so_stance_, EMG_ta_stance_, EMG_pl_stance_] served as the independent variables. For every stride EMG_so_stance_, EMG_ta_stance_ and EMG_pl_stance_ were determined as the median EMG activity during early stance (from heel strike to mid-stance, 0-30% of the stride cycle) multiplied by the duration of this episode in seconds (This is a deviation from the preregistered plans. See “Deviations from the preregistered plans”, II) and demeaned prior to regression. Error_m_ denotes the residual of Model 3.

The foot placement model was created conform Wang and Srinivasan ^13^:

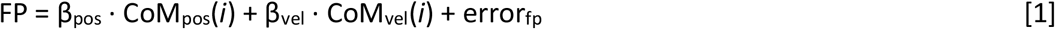

CoM_pos_ and CoM_vel_ were the input for each sample of the step cycle (*i*). Values of error_*fp*_ for CoM state predictors at mid-swing (*i* = 25) and heel strike (*i*=51) were used as the inputs for the ankle strategy model:

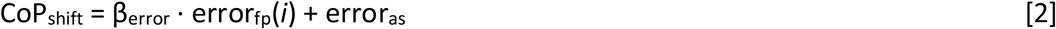

To infer a relationship between the muscle activations and subsequent mediolateral CoP displacement the following regression was performed:

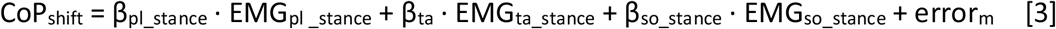

We considered the absolute explained variance of model (2) as a measure of the contribution of the ankle strategy to mediolateral gait stability and the absolute explained variance of model (3) as a measure of the active control underlying this strategy.

### Statistics

All statistical tests were performed in JASP.

### The ankle strategy in steady-state walking

The regression coefficients (β) of model (2) were tested against zero via a Bayesian one-sample t-test, to infer whether mediolateral CoP displacement can be predicted by preceding foot placement error during steady-state walking (H1a) (This is a deviation from the preregistered plans. See “Deviations from the preregistered plans”, III). Similarly, the regression coefficients of model (3) were tested against zero using a Bayesian one-sample t-test to identify whether or not mediolateral CoP displacement correlates with ankle in-/everter muscle activity in the stance phase (H1b). Peroneus longus muscle activity would exert an eversion moment leading to medial CoP displacement, defined by a negative correlation. Tibialis anterior and soleus muscle activity would assist an inversion moment leading to lateral CoP displacement, defined by a positive correlation.

Bayesian equivalents of 2×2 repeated measures ANOVAs with factors Condition (levels: ankle moment constrained/foot placement constrained versus steady-state walking) and Speed (levels: normal versus slow) were used to test the effects of the constraints and walking speed, as well as their interaction, on the contribution of the ankle strategy (see below).

### The ankle strategy with ankle moment constraints

In order to infer whether constraining the ankle strategy was reflected in our ankle strategy model, the absolute explained variances of models (2) and (3) in the ankle strategy constrained condition were tested against the steady-state walking condition. By this, we tested the hypothesis that the contribution of the ankle strategy, in compensating for errors in foot placement, will decrease (i.e. a lower absolute explained variance) when constraining ankle moments (H2).

### The ankle strategy with foot placement constraints

The absolute explained variance of model (2) in the foot placement constrained condition was tested against the steady-state walking condition. This allowed to reveal whether execution of the ankle strategy compensated for constrained foot placement (i.e. a higher absolute explained variance, H3a). The absolute explained variance of model (3) in the foot placement constrained condition was tested against the steady-state walking condition, to evaluate whether this led to compensatory muscle activity (i.e. a higher absolute explained variance, H3b).

### The ankle strategy at different speeds

The absolute explained variances of models (2) and (3) at normal walking speed were tested against the absolute explained variances at slow walking speed. This served to infer whether at a slow walking speed there is a greater contribution of the (active) ankle strategy (i.e. higher absolute explained variances, H4a&b).

### Deviations from the preregistered protocol

A preregistration of the current study’s protocol can be found on OSF: https://osf.io/74pn5. Deviations from the preregistration are described below.

I. We deviated from the preregistered plans, in which we intended to define the mediolateral CoP shift in the global coordinate system. A mediolateral shift in the global coordinate system, would show the CoP displacement perpendicular to the walking direction. However, as we hypothesized an active ankle strategy, we focused on muscles working around the anatomical axes of the feet. Consequently, we realized that our definition of the CoP shift should be based on the effect of these muscles. Inversion and eversion by activating the muscles leads to a mediolateral CoP shift in the local coordinate system of the foot. This would only be the same CoP shift in global coordinates if the local coordinate system would be aligned with the global coordinate system (i.e. if the forward anatomical axis perfectly would be aligned with the walking direction). Participants may demonstrate a different orientation (i.e. pointing the foot more inward or outward), causing a discrepancy between the global mediolateral CoP displacement and the CoP displacement due to ankle moment control. Therefore, the decision to express the CoP shift in the foot’s local coordinate system allows for better interpretation and isolation of the ankle strategy from other strategies such as toeing-out ^20^.
II. We deviated from the preregistered plans, in which we intended to take the integral of the EMG signal over the same time period. To avoid amplification of artefacts by taking the integral, while still retaining the influence of time, we deviated from the preregistered plans and computed the product of the median and time in seconds. (as in van Leeuwen, et al. ^12^)
III. In the preregistered plans, we planned to bootstrap the regression coefficients for each participant. However, although most participants, but not all, demonstrated significant relationships, the final conclusion was based on statistics on group level (testing the regression coefficients against zero in a one-sample Bayesian t-test) and included in the results. (as in van Leeuwen, et al. ^12^)

## Results

### The ankle strategy in steady-state walking

#### Model 2: Ankle strategy model

In the steady-state walking condition (unconstrained), we found extreme evidence (BF_10_ >100) when testing the regression coefficients for error_fp_midswing_ and error_fp_terminalswing_ against zero, for each speed. These findings support that the CoP shift during stance is associated with the error in coordination of foot placement with respect to the CoM (H1). The sign of the relationship was as expected, as it indicates a negative relationship (Figs 3-4). When stepping too medial, participants shifted the CoP more lateral during single stance. When stepping too lateral, participants shifted the CoP more medial during single stance.

**Fig 3.**
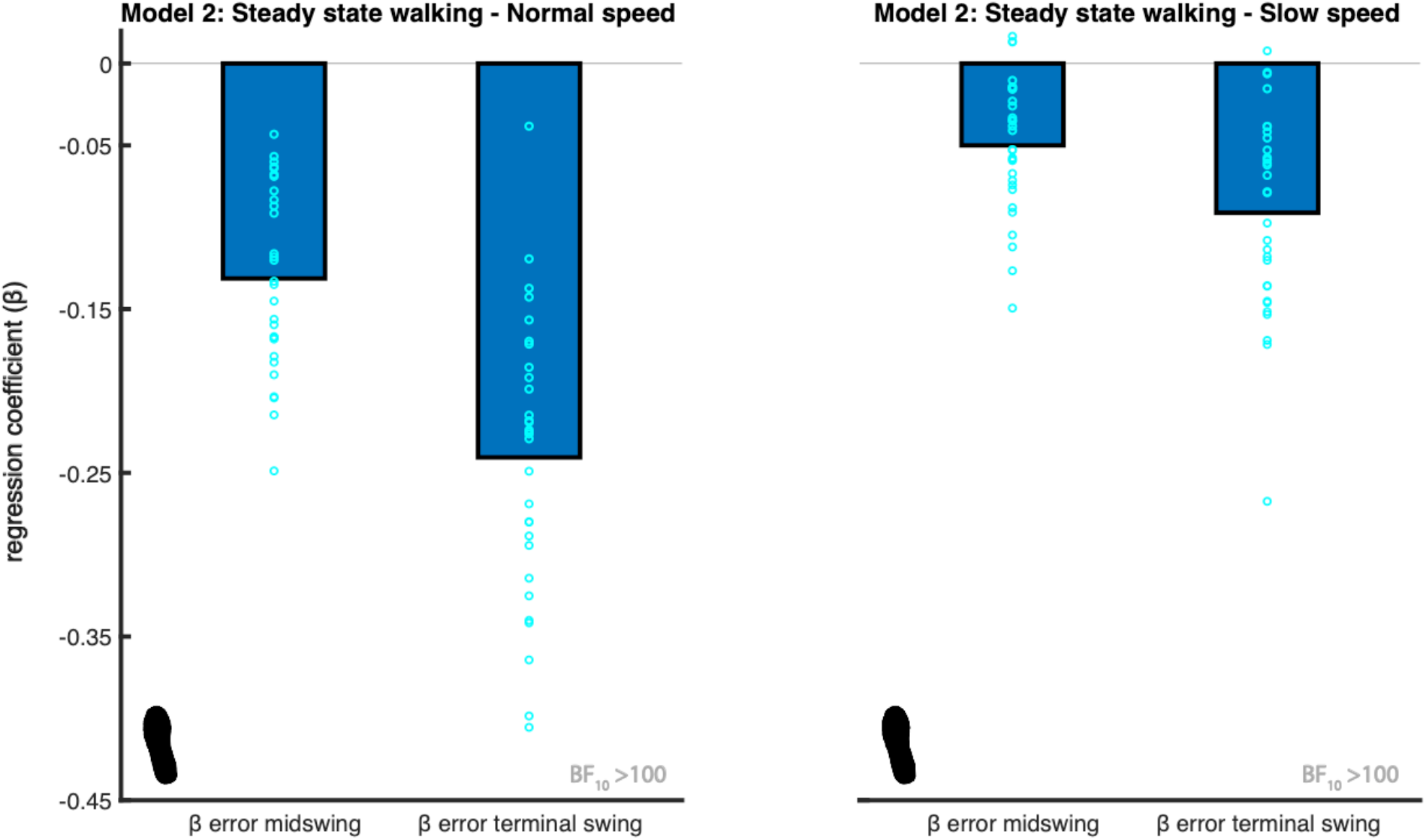
Mean regression coefficients of the ankle strategy model (2). The predictors (β) are the foot placement error at mid-swing and terminal swing. The foot print represents the steady-state walking condition (see Table 1). The light blue circles represent individual data points. The negative relationship shows ankle moments accommodate for stepping inaccuracies, by shifting the CoP in the opposite direction. The Bayes factors (BF_10_) represent the degree of evidence supporting the regression coefficients to be different from zero. This figure was created using Matlab 2021a (https://www.mathworks.com/products/matlab.html) and Adobe Illustrator CC 2018 (https://www.adobe.com/nl/products/illustrator.html).

**Fig 4.**
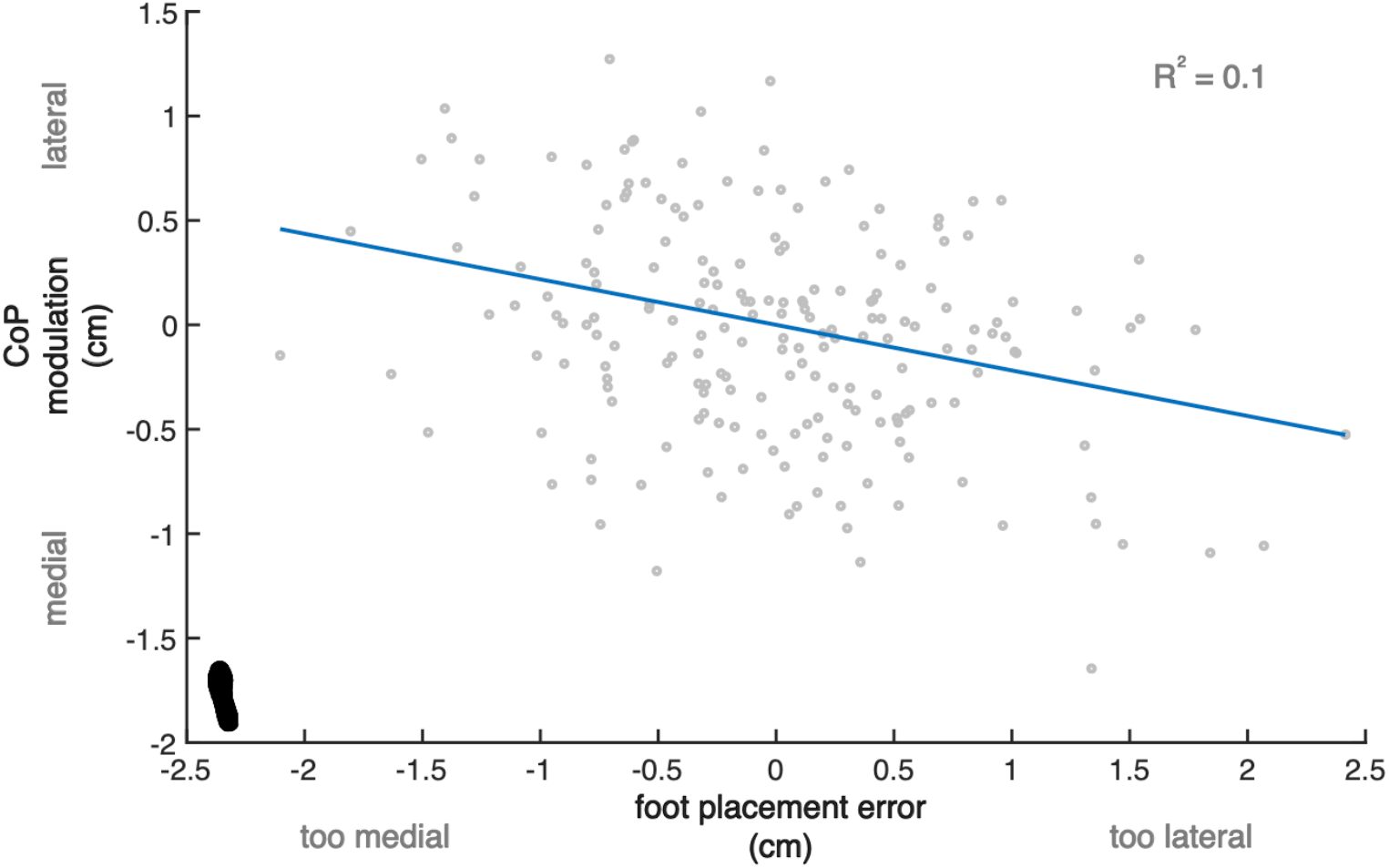
Relationship between foot placement error and subsequent CoP shift. Example of one participant, demonstrating a negative relationship between CoP modulation (ankle moment control) and foot placement error. The blue line represents the fitted model (2) and the data points represent individual (right and left) steps. The foot print represents the steady-state walking condition (see Table 1). Compared to the other participants, this participant demonstrated a relatively high relative explained variance (R^2^ = 0.1). This figure was created using Matlab 2021a (https://www.mathworks.com/products/matlab.html) and Adobe Illustrator CC 2018 (https://www.adobe.com/nl/products/illustrator.html).

#### Model 3: Muscle model

We found extreme evidence (BF_10_ > 100) that the PL coefficient is different from zero at both walking speeds and, as expected, the coefficient was negative at both speeds (Fig 5).

**Fig 5.**
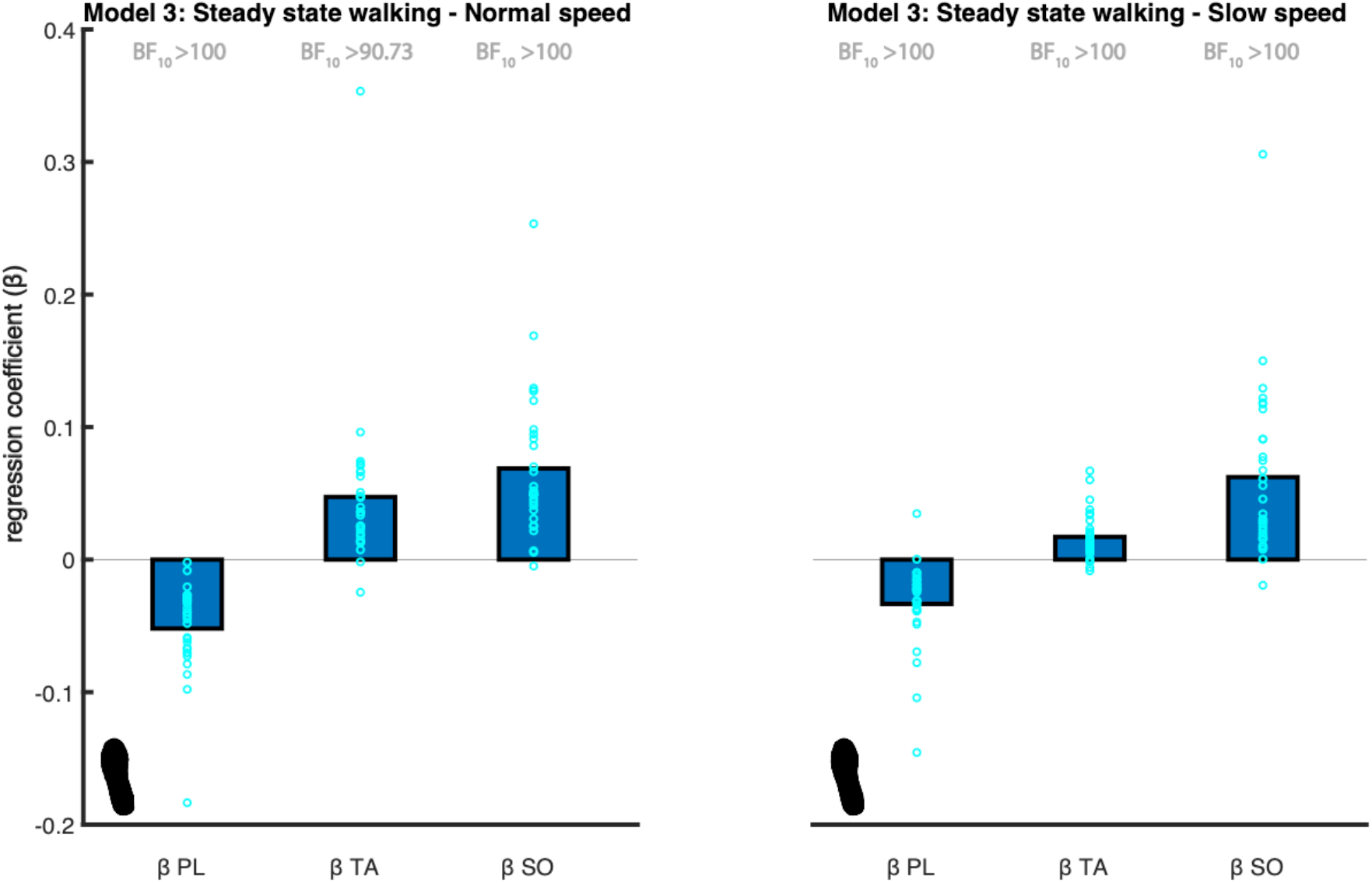
Mean regression coefficients of the muscle model (3). The predictors (β) are the peroneus longus (PL), tibialis anterior (TA) and soleus (SO) muscles’ activity. The foot print represents the steady-state walking condition (see Table 1). The light blue circles represent individual data points. The Bayes factors (BF_10_) represent the degree of evidence supporting the regression coefficients to be different from zero. This figure was created using Matlab 2021a (https://www.mathworks.com/products/matlab.html) and Adobe Illustrator CC 2018 (https://www.adobe.com/nl/products/illustrator.html).

We found very strong evidence (BF_10_ = 90.733) that the TA coefficient is different from zero at a normal walking speed and extreme evidence (BF_10_ > 100) at a slow walking speed. As expected, the coefficient was positive at both speeds (Fig 5).

The soleus muscle assists inversion leading to lateral CoP displacement. As such we expected a positive association. We found extreme evidence (BF_10_ > 100) supporting the SO coefficient to be different from zero at both walking speeds. As expected, the coefficient was positive at both speeds (Fig 5).

Combined the results support an active contribution to ankle moment control (H2).

#### The ankle strategy with ankle moment constraints

Bayesian repeated-measures ANOVA of the absolute explained variance of the ankle strategy model (2), including the ankle moments constrained and the steady-state walking condition at both speeds, demonstrated that the best model included Condition, Speed and their interaction, for the error in foot placement predicted at both mid-swing and terminal swing (Fig 6). We found extreme evidence for this model as compared to the null model (BF_10_mid_ = 3.091·10^17^, BF_10_ts_ =1.283·10^20^).

**Fig 6.**
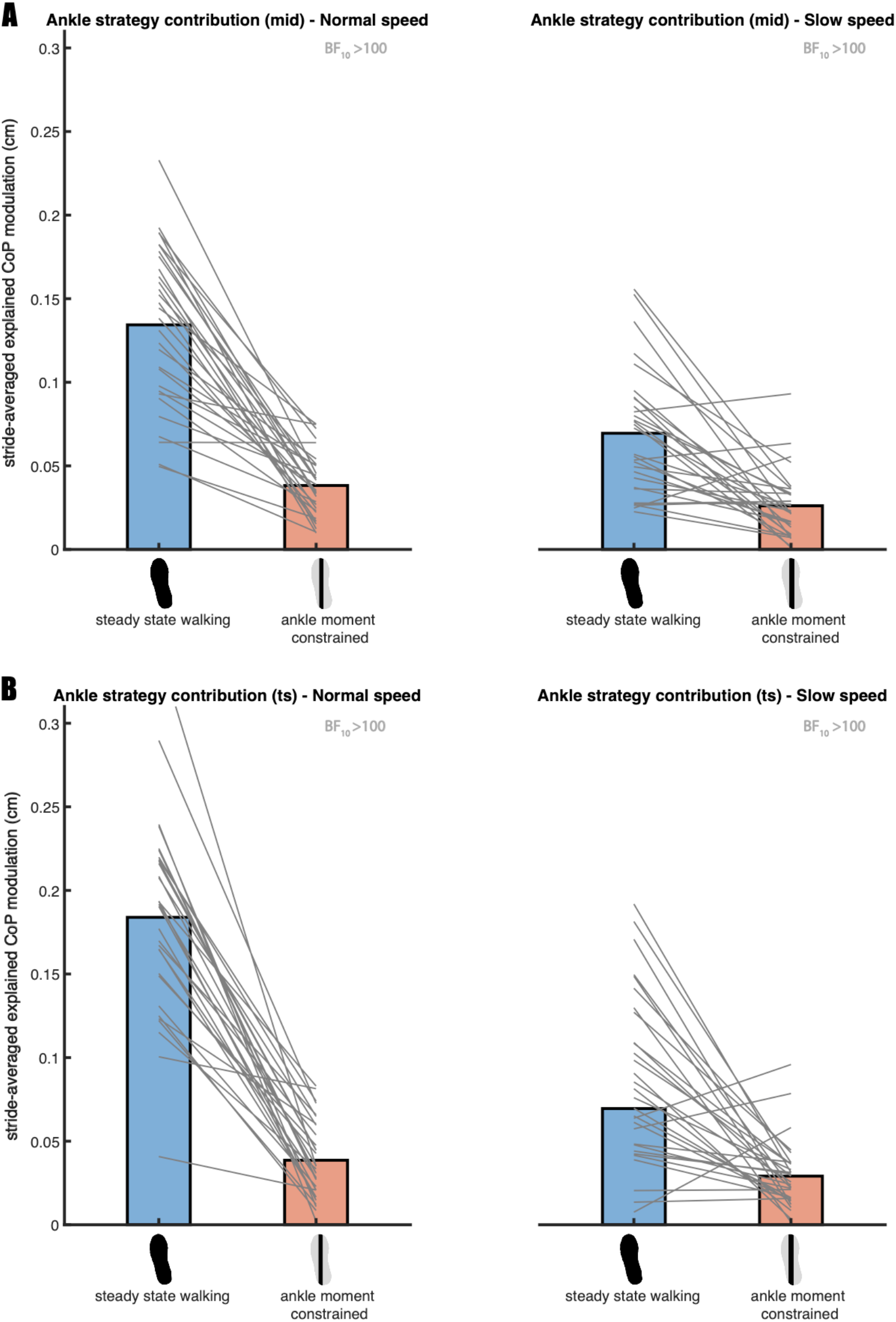
Mean absolute explained variance of the ankle strategy model (2). For normal (left) and slow (right) walking speeds panels A and B depict respectively the absolute explained variances for the foot placement error at mid- and terminal swing. Blue and red bars represent the steady-state walking and ankle moment constrained conditions respectively. In this figure the absolute explained variance is expressed as the square root of the stride averaged explained variance, as a reflection of the magnitude of the average explained CoP shift in centimeters. The grey lines connect individual data points. The Bayes factors (BF_10_) denote the degree of evidence for a decreased absolute explained variance in the ankle moment constrained condition, as compared to steady-state walking. This figure was created using Matlab 2021a (https://www.mathworks.com/products/matlab.html) and Adobe Illustrator CC 2018 (https://www.adobe.com/nl/products/illustrator.html).

Given the interaction, we conducted a post hoc one-tailed Bayesian paired samples t-test for each speed independently. At normal walking speed (Fig 6), we found extreme evidence supporting a decreased absolute explained variance (BF_10_mid_ = 2.736·10^6^, BF_10_ts_ = 7.471·10^6^) in the ankle moment constrained as compared to the steady-state walking condition. At slow walking speed (Fig 6), extreme evidence supported a decreased absolute explained variance as well (BF_10_mid_ =278.158, BF_10_ts_ = 577.083). Combined these results indicate that the LesSchuh effectively constrained ankle moment control to correct for foot placement errors (H2).

### The ankle strategy with foot placement constraints

Bayesian repeated measures ANOVA of the absolute explained variance of the ankle strategy model (2), including the foot placement constrained and the steady-state walking conditions at both speeds, demonstrated that the best model included only Speed as a factor (Fig 7). We found extreme evidence for this model as compared to the null model (BF_10_mid_ = 3.510·10^9^, BF_10_ts_ = 9.158·10^14^). Since the factor Condition was not included in the best model, we found no evidence that the contribution of the ankle strategy increases when foot placement is constrained (H3a). Consequently, there is no evidence for compensatory muscle activity related to the ankle strategy (H3b).

**Fig 7.**
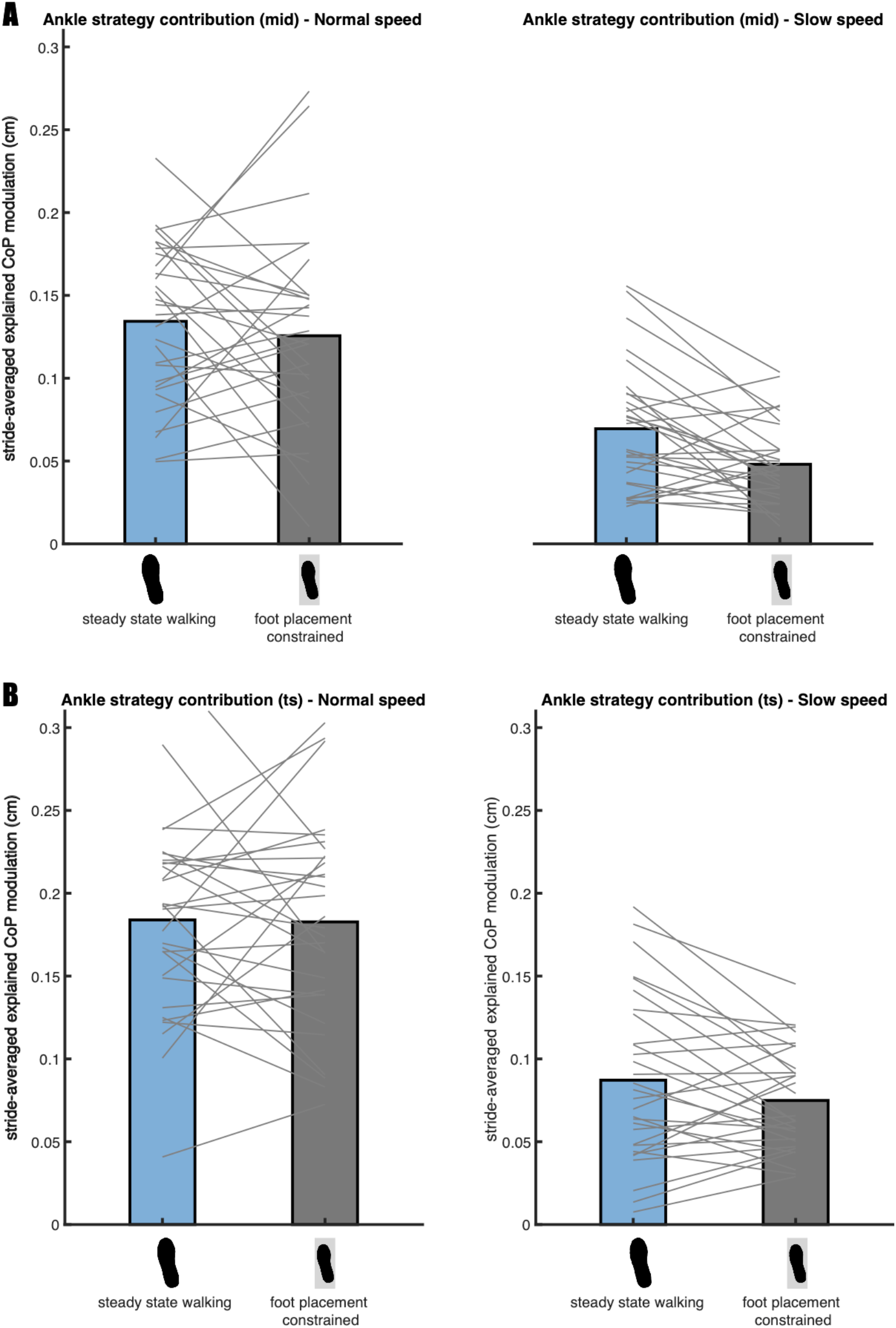
Mean absolute explained variance of the ankle strategy model (2). For normal (left) and slow (right) walking speeds panels A and B depict respectively the absolute explained variances for the foot placement error at mid- and terminal swing. Blue and grey bars represent the steady-state walking and foot placement constrained conditions respectively. In this figure the absolute explained variance is expressed as the square root of the stride averaged explained variance, as a reflection of the magnitude of the average explained CoP shift in centimeters. The grey lines connect individual data points. As the best model did not include the factor Condition, no Bayes factors have been presented. This figure was created using Matlab 2021a (https://www.mathworks.com/products/matlab.html) and Adobe Illustrator CC 2018 (https://www.adobe.com/nl/products/illustrator.html).

### The ankle strategy at different speeds

We did not find any evidence that the (active) contribution of the ankle strategy is higher at a lower speed (H4a&B).

The Bayesian repeated measures ANOVAs of the absolute explained variance of the ankle strategy model (2), included the factor Speed. We tested the absolute explained variance during steady-state walking at normal and slow walking speed against each other in a one-tailed Bayesian paired samples t-test. We found very strong evidence (BF_10_mid_ = 0.022, BF_10_ts_ = 0.017) against a higher absolute explained variance during slow walking (H4a). Consequently, there is no evidence for muscle activity associated to a higher ankle strategy contribution at a slow speed (H4b).

Visual inspection of figures 6 and 7 indicated a trend opposite to what we initially expected. The absolute explained variance of the ankle strategy model (2) appeared higher at a normal as compared to at a slow walking speed. Hence, we performed an exploratory one-tailed Bayesian paired samples t-test and found extreme evidence (BF_10_mid_ = 3034.947, BF_10_ts_ = 62158.215) that the contribution of the ankle strategy, conform model (2), is indeed higher at a normal speed.

When performing a similar exploratory one-tailed Bayesian paired samples t-test for the muscle model’s (3) absolute explained variance between speeds, extreme evidence (BF_10_ = 436273.091) supported a higher active contribution to execution of the ankle strategy at normal as compared to slow walking speed (Fig 8).

**Fig 8.**
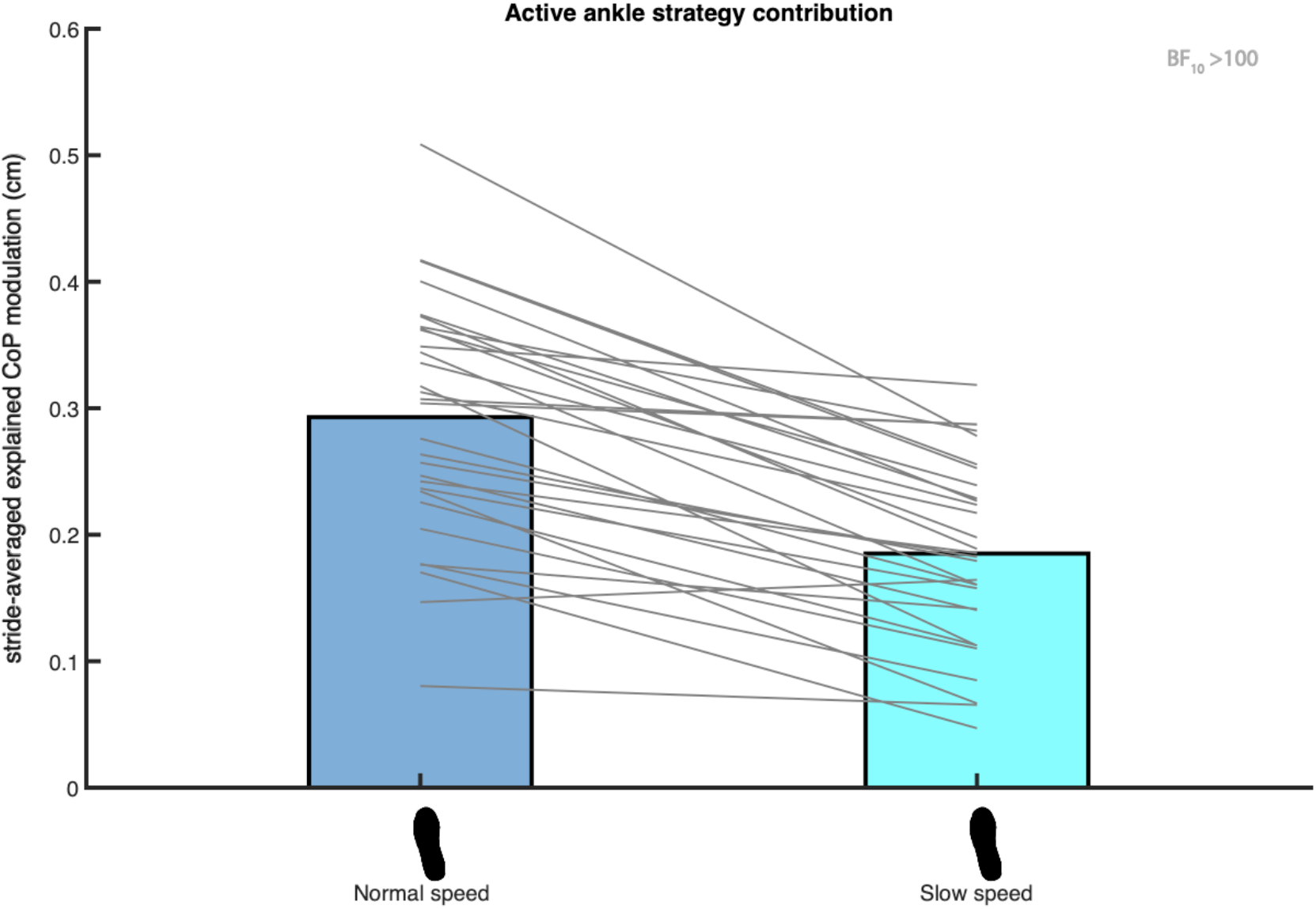
Mean absolute explained variance of the muscle model (3). Blue and light blue bars represent respectively the absolute explained variance at normal and slow walking speed. In this figure the absolute explained variance is expressed as the square root of the stride averaged explained variance, as a reflection of the magnitude of the average explained CoP shift in centimeters. The foot print represents the steady-state walking condition (see Table 1). The grey lines connect individual data points. The Bayes factor (BF_10_) denotes the degree of evidence for a lower absolute explained variance at slow as compared to at normal walking speed. This figure was created using Matlab 2021a (https://www.mathworks.com/products/matlab.html) and Adobe Illustrator CC 2018 (https://www.adobe.com/nl/products/illustrator.html).

## Discussion

In a previous study, we investigated foot placement control with respect to CoM state, as a stability control strategy ^12^. In the current study, we investigated whether inaccurate execution of this strategy may be compensated for by performing an (active) ankle strategy. During steady-state walking, we demonstrated a relationship between errors in foot placement and subsequent mediolateral CoP displacement during single stance. The here-reported negative relationship indicates that steps that are too medial are compensated for by a lateral CoP shift and vice versa that steps that are too lateral are compensated for by a medial CoP shift. It seems that inaccuracies in foot placement, potentially leading to instability, can be mitigated by a mediolateral ankle strategy. Ankle strategy CoP shifts were associated with muscle activity around the ankle, indicating an active implementation of this strategy during single stance. Ankle moment control was effectively constrained by LesSchuh, but unexpectedly, when walking with constrained foot placement, ankle moment control did not increase as a compensation.

### Compensatory ankle strategy

During unconstrained steady-state walking, ankle moments contribute to CoP control after foot placement. Walking with LesSchuh (Fig 1) served to support this. Concurrent with a reduction in the absolute explained variance of the ankle strategy model (Fig 6), compensatory increases in step width and frequency were adopted, arguably to maintain a stable gait pattern ^12^. During steady-state walking, the CoP shifts can be predicted by the error of the foot placement model. This underscores that the foot placement model represents a stability control strategy, which is complemented by a mediolateral ankle strategy. The complementary nature of the foot placement and ankle strategies is in line with earlier findings in the literature, mostly on responses to perturbations, but also on steady-state walking ^7-9,11,27^. However, we found a low R^2^ (S2 Fig 2) and absolute explained variance (Fig 6-8). Given the extreme evidence for inclusion of the predictors in the model (Fig 3), we conjecture that step-by-step modulations in single stance CoP displacement can be explained by the errors in foot placement, albeit only partially. Below we consider explanations as to why this relationship explains only part of the variance in mediolateral CoP displacement.

Firstly, we explored the possibility that only steps that are too medial require execution of an (active) ankle strategy in S2. We suggested that stepping too lateral does not pose a direct threat to stability, whereas stepping too medial could lead to a sideward fall. Yet, based on our exploratory analysis (S2), there was no evidence that including only those steps that were too medial in the regression improved the predictability of the model predictions. Apparently, a mediolateral ankle strategy is used equally to compensate for steps that are too medial and steps that are too lateral. This is in line with Brough, et al. ^27^, who demonstrated significant ankle inversion moment responses to both medial and lateral foot placement perturbations, despite smaller effects on dynamic balance following lateral foot placement perturbations.

Secondly, our definition of the mediolateral ankle strategy is limited to single stance only. However, between inaccurate foot placement and single stance ankle moment control, other stability control strategies can already compensate for inaccurate foot placement. For example, push-off can directly influence the CoM trajectory ^11,19,28,29^, mitigating the need for a compensatory CoP shift during single stance. Furthermore, during the double support phase, the CoP could already be modulated by an ankle moment ^27^. CoP displacement during double stance has been shown to control CoM velocity in response to perturbations ^30^. Given that in our study ankle moment control did not compensate for constrained foot placement (Fig 7), it is likely that other strategies and/or ankle moment control during double stance played a role in maintaining stability despite inaccurate foot placement.

Thirdly, foot placement error might not cause instability even if only partially compensated for. As such, the ankle moments do not need to be and are not tightly controlled.

A different temporal order in which the foot placement and ankle strategy complement each other offers a potential fourth explanation for our low R^2^ value. Our ankle strategy model (model (2)) describes complementary CoP shifts to compensate for inaccurate foot placement. This is in line with the ankle strategy as a corrective mechanism for inaccurate foot placement ^8^. However, the foot placement-ankle moment relationship can also be considered in the reversed temporal order, with ankle moment control preceding foot placement control, more like an early response to a perturbation ^7^. Evaluating the latter temporal order, Fettrow, et al. ^11^ demonstrated an inverse relationship between the contributions of the foot placement and ankle strategy in response to visual perturbations. They found a similar inverse relationship between foot placement and ankle strategy contributions during steady-state walking.

Given Fettrow, et al. ^11^’s considerably higher R^2^ in explaining the relationship between foot placement and ankle moment control, it could be that the CoP control through ankle moments is more important preceding foot placement, rather than as a compensation following foot placement. There are two interpretations of the coupling between the ankle moment and foot placement control, given that ankle moment control precedes foot placement. The first interpretation suggests that foot placement corrects for inaccuracies in preceding ankle moments ^11^. The second interpretation, based on perturbed targeted stepping, proposes that ankle moment control assists subsequent foot placement ^31^. The latter interpretation would imply that participants walking with LesSchuh stepped less accurately according to the foot placement model, as a result of decreased assistance by ankle moment control ^12^. It seems that execution of the ankle and foot placement strategy cannot be considered separately, because of their inherent interaction.

Admittedly, there are some limitations in the way we expressed the CoP shift during single stance. We determined the CoP shift based on an average value with respect to the initial single stance CoP position, similar to Hof, et al. ^2^. While averaging typically reduces noise-induced prediction errors, it may remove information that could enhance predictability. Furthermore, we expressed the CoP in the local coordinate system of the foot, irrespective of its position relative to the CoM, unlike others ^11,15^. Although our foot placement error provides a relationship with the CoP based on CoM state, this choice reduces CoM position and speed to a unidimensional value. In doing so we inevitably lost information, although exploring inclusion of CoM speed at terminal swing did not improve the ankle strategy model’s predictability (S3). Another source of information that we are missing is single stance time, which earlier studies ^11,15^ incorporated by taking the integral of the distance between the CoM and the CoP during single stance. Since we asked participants to walk according to a metronome beat, this would most likely only have affected our comparison between speeds.

### Speed effect

Differences in CoP shift calculations could have contributed to the unexpected results regarding the contribution of the ankle strategy at different speeds. Based on the absolute explained variance we observed a higher ankle strategy contribution at normal as compared to slow walking speed during steady-state walking (Fig 8). In contrast, Fettrow, et al. ^15^ demonstrated that, in response to perturbations, the CoP shifted more prominently at a slower speed as compared to at a faster walking speed, with the difference emerging predominantly during the double support phase. The discrepancy with our results appears to suggest that speed effects in perturbation studies ^15^ cannot be generalized directly to steady-state walking. Perhaps, ankle moment control is less important at a slow steady-state walking speed, with more scope for an additional shift in case of a perturbation, as compared to at a faster walking speed.

During steady-state walking, foot placement is more tightly controlled at faster speeds ^14^. If execution of the ankle strategy mainly serves to accurately control subsequent foot placement, one would thus expect a larger contribution of the ankle strategy at normal compared to slow walking speeds. Along this line of reasoning, the visually observed larger detrimental effect of LesSchuh on the degree of foot placement control at normal as compared to slow walking speed ^12^ could be explained. However, if execution of the ankle strategy mainly serves to attenuate errors after foot placement, one would expect a higher contribution of the ankle strategy at slow walking speeds, based on the lower degree of foot placement control at slower speeds ^14^. That is, if the lower contribution of the ankle strategy during slow walking does not reflect a general lower demand for tight stability control at slower speeds. Further research is needed to clarify the most important role of the mediolateral ankle strategy during unconstrained steady-state walking.

### Passive or active?

We identified an active mediolateral ankle strategy. Previous studies demonstrated ankle muscle responses following perturbations ^7,15^. We show that also during steady-state walking peroneus longus, tibialis anterior and soleus muscle activity during single stance are associated with the subsequent CoP shift (Fig 4). Yet, it should be noted that R^2^ values were low (S3). This can be due to our assumption of a linear relationship being too simple, but also suggests a contribution of passive dynamics during single stance.

## Conclusion

We sought to investigate the mediolateral ankle strategy as a compensatory mechanism for inaccurate foot placement. Confirming such a compensatory mechanism, we identified a relationship between errors in foot placement and subsequent CoP shifts during single stance. Furthermore, correlations between ankle muscle activity and mediolateral CoP shifts provide evidence for active control. However, our findings do not encompass the entire complementary nature of the foot placement and ankle strategy. Foot placement and ankle moment control are ‘caught’ in a circular relationship. Given diminished foot placement control when ankle moments are constrained, there seems to be a direct contribution of ankle moment control to foot placement. One can hence not fully identify the control mechanism by considering the two strategies separately nor by considering them in a fixed temporal order. Likely, shifting the CoP is not only aimed at “fixing” the current step but also on enhancing control of subsequent steps. When ankle moments are constrained, one does not step more accurately and vice versa when foot placement is constrained, ankle moment control does not act as a compensatory mechanism. During steady-state walking muscle activity drives both foot placement and ankle moment control, and it follows that by constraining one, one also constrains the other.

## Acknowledgements

The authors are thankful for all participants, technical support and assistance during the data collection, especially for Leon Schutte, who built LesSchuh. Sjoerd Bruijn and Moira van Leeuwen were funded by a grant from the Netherlands Organization for Scientific Research (016.Vidi.178.014), https://www.nwo.nl/en/.

## Authors’ contributions

MvL performed the experiments. All authors contributed to the experimental design and the writing/editing of the manuscript.

## Supplementary material

### S1

We instructed participants to keep placing their feet in a similar orientation (straight ahead) throughout all conditions, to avoid a toeing out strategy, especially in the ankle moment constrained condition, where we wanted to limit the CoP shift during single stance. Following Fig S1, showing the toe-out angle of the 30 included participants, it appears most participants followed these instructions. The toe-out angle (with respect to the global vertical axis) showed to be similar during steady-state and ankle moment constrained walking, given moderate evidence against any difference at both normal (BF_10_ =0.319) and slow (BF_10_ = 0.196) speed.

**S1 Fig 1.**
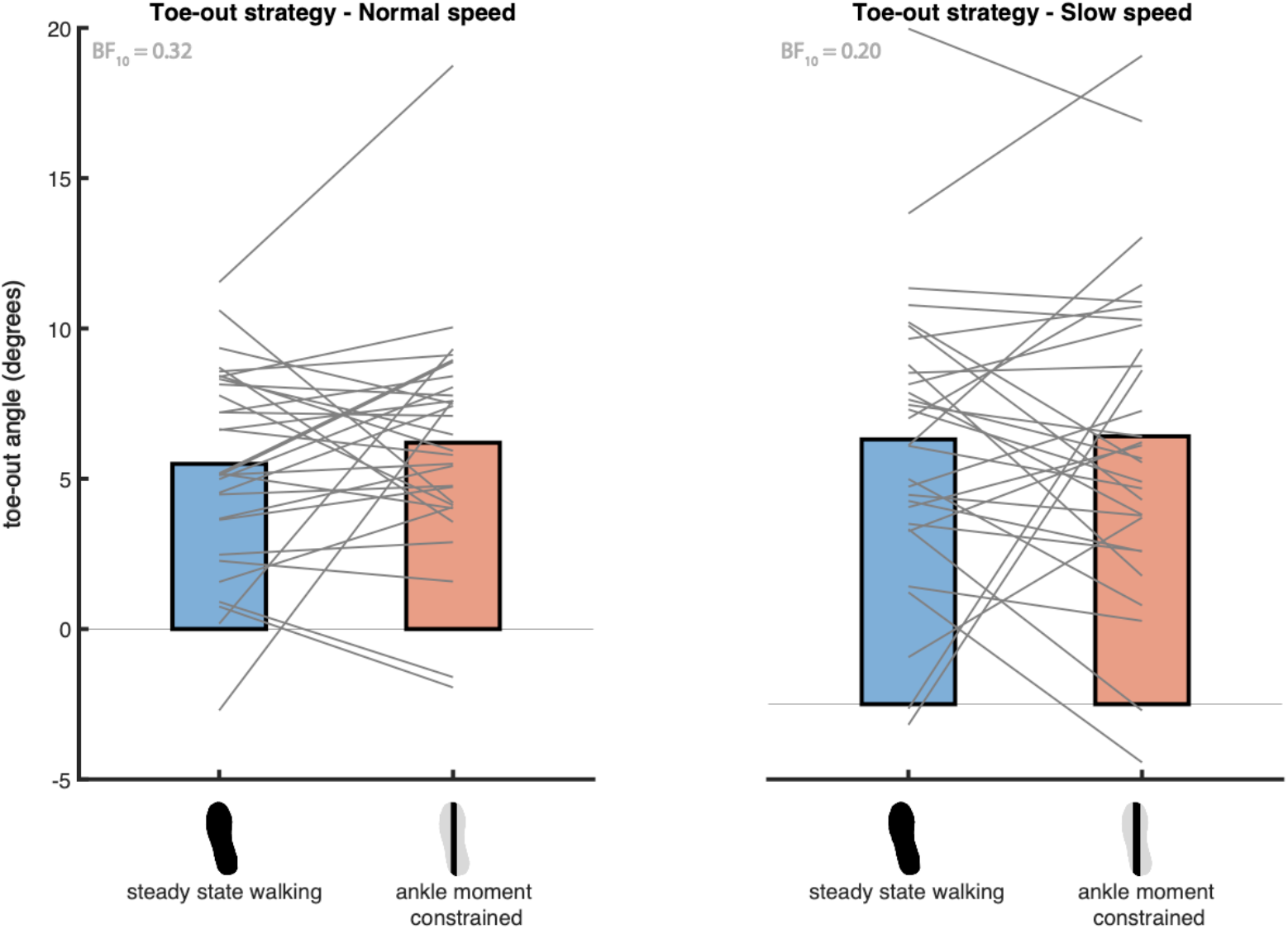
The toe-out angle during steady-state and ankle moment constrained walking. The angles are expressed with respect to the feet pointing straight ahead (0 degrees). Positive and negative angles correspond to respectively toeing-out and toeing-in. The grey lines connect individual data points. The Bayes factors (BF_10_) represent the degree of evidence against an increased toe-out angle in the ankle moment constrained condition, as compared to steady-state walking. This figure was created using Matlab 2021a (https://www.mathworks.com/products/matlab.html) and Adobe Illustrator CC 2018 (https://www.adobe.com/nl/products/illustrator.html).

### S2

#### Only including medial steps for the ankle strategy model

One could argue, that stepping too lateral does not threaten stability like stepping too medial does. Stepping too medial can lead to a sideward fall, whereas a “fall” induced by stepping too lateral would merely accelerate the body towards the subsequent stance foot. Therefore, we also considered the ankle strategy model while only including steps which were too medial.

The negative relationship between foot placement error and the CoP shift was retained in the model including only medial steps. When testing the regression coefficients against zero using a Bayesian one-sample t-test, extreme evidence was found for the ankle strategy models at both speeds.

**S2 Fig 1.**
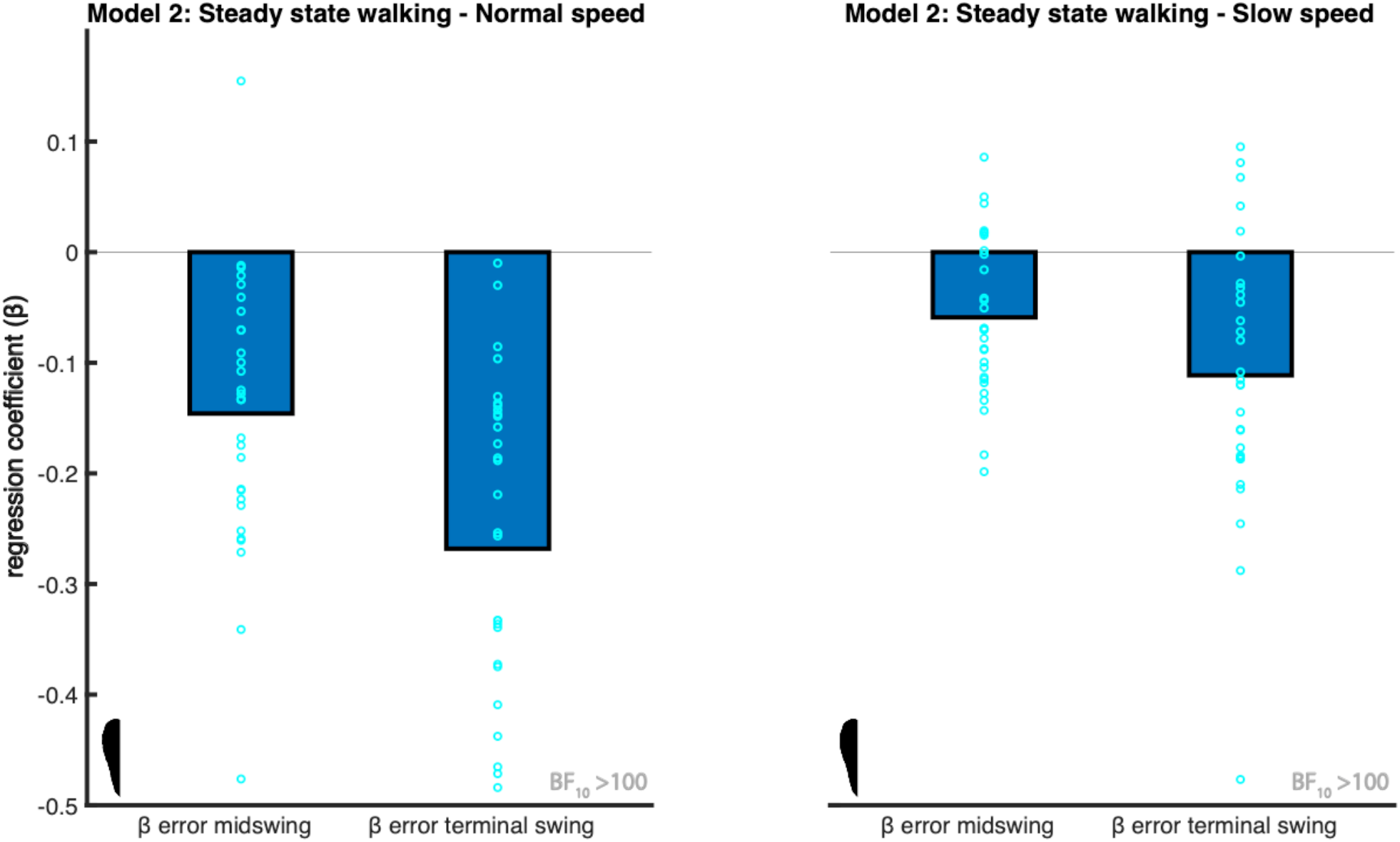
Mean regression coefficients of the ankle strategy model (2). The predictors (β) are foot placement error at mid-swing and terminal swing. In contrast to Fig 3, here only the foot placement errors corresponding to the too medial steps were included, as indicated by half a steady-state walking condition foot print. The light blue circles represent individual data points. The negative relationship shows ankle moments accommodate for stepping inaccuracies, by shifting the CoP more lateral when stepping more medial. The Bayes factors (BF_10_) denote the degree of evidence supporting the regression coefficients to be different from zero. This figure was created using Matlab 2021a (https://www.mathworks.com/products/matlab.html) and Adobe Illustrator CC 2018 (https://www.adobe.com/nl/products/illustrator.html).

S2 Fig 2a shows that the relative explained variance appears to be higher when including all steps in the model as compared to including only too medial steps, with foot placement error at mid-swing as predictor. Based on the mid-swing foot placement error this was supported by very strong evidence at normal walking speed (BF_10_ = 79.173). Yet, at a slow walking speed there is anecdotal evidence against this observation (BF_10_ = 0.683). Similarly, S2 Fig 2b shows a trend for a higher relative explained variance when all steps are included in the model with foot placement error at terminal swing as predictor. Testing this observation (two-tailed), we found extreme evidence supporting this observation at normal speed (BF_10_ = 990100.061) and anecdotal evidence against this observation at slow speed (BF_10_ = 0.476).

**S2 Fig 2.**
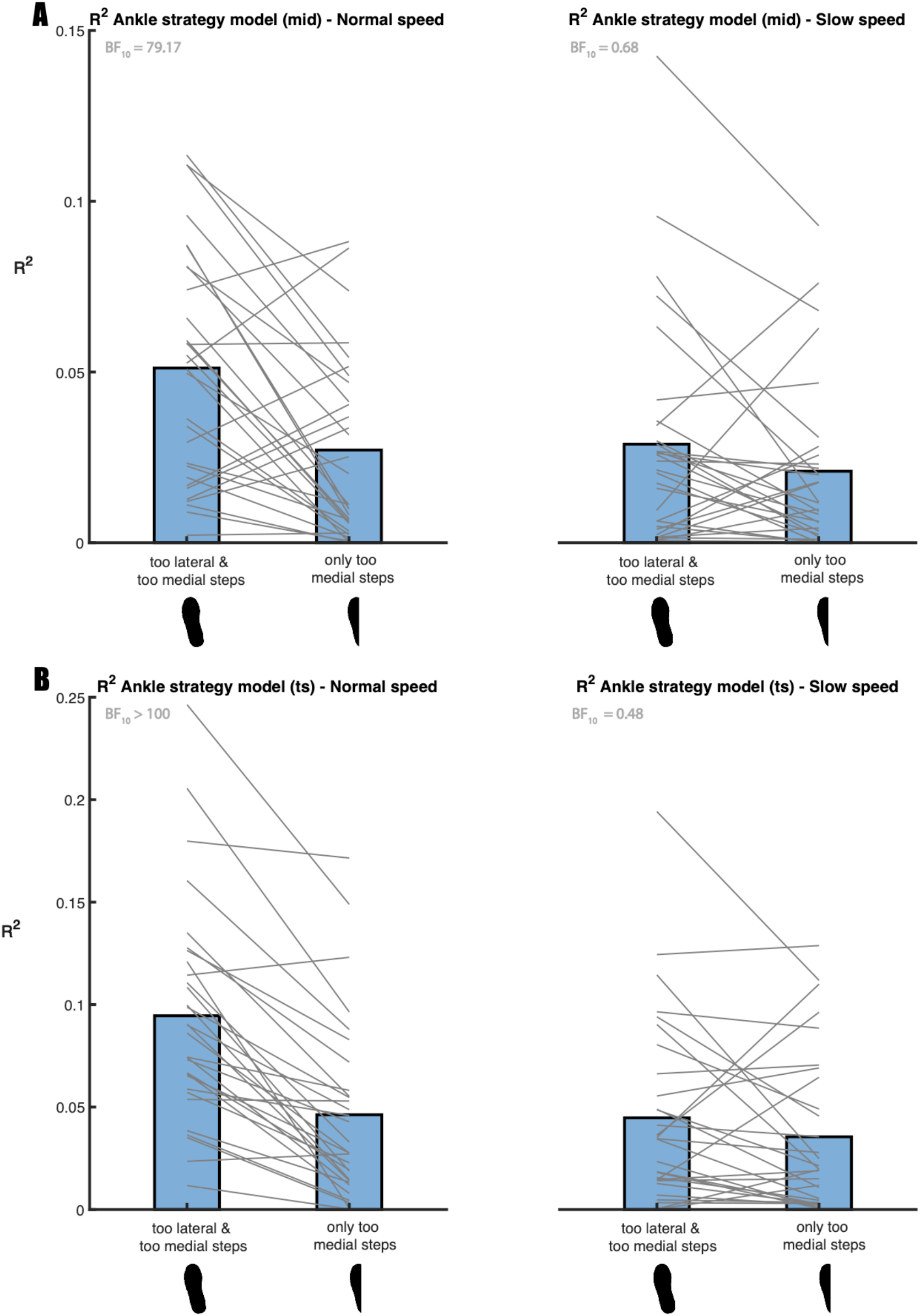
The relative explained variance (R^2^) of the ankle strategy model (2). R^2^ was computed for the model with the foot placement error at mid-swing (panel A) and terminal swing (panel B). A comparison is made between including all (full steady-state walking condition foot print) as compared to only too medial steps (half a steady-state walking condition foot print), with the foot placement error at mid-swing as predictor. For most participants, the ankle strategy model, explained a higher percentage of the variance in CoP shifts, when both the too lateral and the too medial steps are included. Grey lines connect individual data points. The Bayes factors (BF_10_) denote the degree of evidence. This figure was created using Matlab 2021a (https://www.mathworks.com/products/matlab.html) and Adobe Illustrator CC 2018 (https://www.adobe.com/nl/products/illustrator.html).

### S3

#### Relative explained variance ankle strategy muscle model (3)

**S3 Fig 1.**
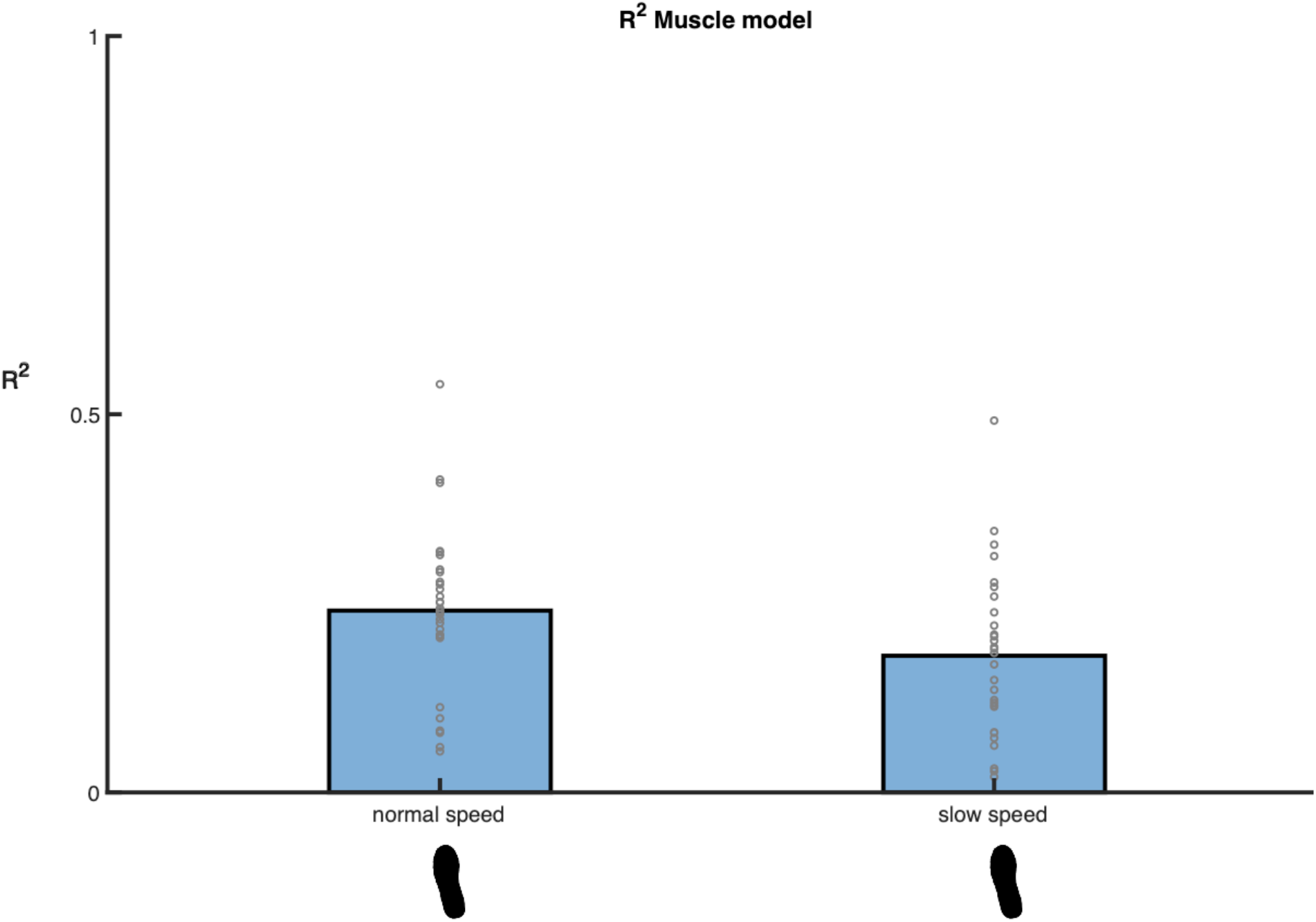
R^2^ of the ankle strategy muscle model (3). The bars represent the average R^2^ at respectively normal (left) and slow (right) walking speed. Grey circles represent individual data points. This figure was created using Matlab 2021a (https://www.mathworks.com/products/matlab.html) and Adobe Illustrator CC 2018 (https://www.adobe.com/nl/products/illustrator.html).

